# Utilization of cobalamin is ubiquitous in early-branching fungal phyla

**DOI:** 10.1101/2020.10.13.337048

**Authors:** Małgorzata Orłowska, Kamil Steczkiewicz, Anna Muszewska

**Affiliations:** Institute of Biochemistry and Biophysics, Polish Academy of Sciences, Pawinskiego 5A, 02-106 Warsaw, Poland

## Abstract

Cobalamin is a cofactor present in essential metabolic pathways in animals and one of the water-soluble vitamins. It is a complex compound synthesized solely by prokaryotes. Cobalamin dependence is scattered across the tree of life. In particular, fungi and plants were deemed devoid of cobalamin. We demonstrate that cobalamin is utilized by all fungal lineages, except for Dikarya. This observation is supported by the genomic presence of both B12 dependent enzymes and cobalamin modifying enzymes. Moreover, the genes identified are actively transcribed in many taxa. Most fungal cobalamin dependent enzymes and cobalamin metabolism proteins are highly similar to their animal homologs. Phylogenetic analyses support a scenario of vertical inheritance of the cobalamin trait with several losses. Cobalamin usage was probably lost in Mucorinae and at the base of Dikarya which groups most of the model organisms which hindered B12-dependent metabolism discovery in fungi. Our results indicate that cobalamin dependence was a widely distributed trait at least in Opisthokonta, across diverse microbial eukaryotes and likely in the LECA.

## Introduction

Cobalamin, also known as vitamin B12, is the most common cobalt-containing compound in nature and one of eight known water-soluble vitamins grouped into B class. Cobalamin is an organometallic complex compound that contains a cobalt atom placed within a corrin ring. Vitamin B12 is derived from uroporphyrinogen III, which is also the first macrocyclic intermediate in a common pathway of heme and chlorophyll biosynthesis (Chatthanawaree 2011)(Dereven’kov et al. 2016)(Frank et al. 2005). In animals, cobalamin is used as a cofactor in myelin formation and thus is crucial for the proper functioning of the nervous system. A deficit of this vitamin in the diet may lead to sensory or motor deficiencies and to degeneration of the spinal cord (Dardiotis et al. 2017).

Biosynthesis of cobalamin takes place only in bacteria and archaea which is quite unique for such a widely used vitamin. It is a very complex process and involves more than 30 genes (Roth et al. 1993) collectively conserved only in B12-producing prokaryotes which suggest a common origin of the whole pathway. Nonetheless, animals and protists utilize cobalamin in their metabolism so they have to intake this vitamin with food. Interestingly, also other eukaryotic microorganisms, including Phytophthora (Oomycota) and Dictyostelium (Amoebozoa), do possess B12-dependent enzymes. Some algae like *Porphyridium purpureum* and *Amphidinium operculatum*, can obtain the cobalamin cofactor from associated bacteria (Croft et al. 2005). Plants and fungi are believed to neither synthesize nor even have a need for the cobalamin (Duda et al. 1957; Jah et al. 2002). Even more, they are regarded as devoid of cobalt at all (Zhang et al. 2019).

In Eukaryotes, B12-dependent enzymes are used in diverse processes ranging from DNA replication and repair, regeneration of methionine from homocysteine, catabolic breakdown of some amino acids into succinyl-CoA (necessary for citric acid cycle) and proper myelin synthesis. Only eight enzymes from the above pathways seem to be uniquely present in B12-dependent organisms. They either modify cobalamin or use it as a cofactor. The former group contains methylmalonyl Co-A mutase-associated GTPase Cob (MeaB), cob(I)yrinic acid a,c-diamide adenosyltransferase (CblAdo transferase), cyanocobalamin reductase (CblC) and cobalamin trafficking protein D (CblD) proteins, while the latter includes ribonucleotide reductase class II (RNR class II), methionine synthase (MetH), methylmalonyl-CoA epimerase (MM-CoA epimerase) and methylmalonyl-CoA mutase (MM-CoA mutase). All these proteins are present in animals, including Holozoa e.g. *Monosiga brevicollis*. For consistency and clarity, we will use the names of human representatives (given above) to tag the above eight enzymes.

RNR is an enzyme that catalyzes the formation of deoxyribonucleotides from ribonucleotides. It plays a pivotal role in synthesis, reparation, and regulation of the total rate of DNA synthesis (Herrick and Sclavi 2007)(Larsson et al. 2004). RNRs are divided into three classes that are working based on similar mechanisms but using a different compound to generate free radicals. Class I reductases are divided into IA and IB subclasses. These reductases generate tyrosyl free radicals from iron. Subclass IA is distributed in eukaryotes, eubacteria, and viruses. Subclass IB can be found only in eubacteria. Class II reductases use free radicals from cobalamin and are distributed in archaebacteria, eubacteria, and bacteriophages. The same distribution applies to class III, but this class uses a glycine radical. Most eukaryotes, including animals, use class IA reductases, but surprisingly *Phytophthora* spp. uses cobalamin-dependent class II RNR.

Methionine synthetase (MetH) comes in two variants: cobalamin-dependent MetH (EC 2.1.1.13) and cobalamin-independent MetE (EC 2.1.1.14). MetH catalyzes the final step in the remethylation of homocysteine which explains increased levels of homocysteine upon vitamin B12 deficiency. In animals, this may lead to blindness, neurological symptoms, and birth defects (Outteryck et al. 2012). MetH requires Cyanocobalamin reductase (CblC) and Cobalamin trafficking protein (CblD) for proper function. ClbC catalyzes the decyanation of cyanocobalamin and the dealkylation of alkylcobalamins. In bacteria, an analog of CblC/D, namely TonB, is involved in energy transduction for the uptake of cobalamin (Lerner-Ellis et al. 2006)(Hannibal et al. 2009). CblD interacts with CblC and directs CblC-cob(II)alamin molecules to the mitochondrion. Consistently, ClbC localizes either to cytoplasm or mitochondria, while ClbD remains in the cytosol (Gherasim et al. 2013; Mah et al. 2013).

CblAdo transferase, cob(I)yrinic acid a,c-diamide adenosyltransferase, converts cobalamin into adenosylcobalamin (AdoCbl) (Mera and Escalante-Semerena 2010)(Marsh and Meléndez 2012) multiple enzymes that catalyze unusual rearrangement or elimination reactions some of them are restricted to prokaryotes e.g. lysine-5,6-aminomutase, isobutyryl-CoA mutase and glutamate mutase other are present also in Eukaryotes eg. methylmalonyl-CoA mutase.

In humans, MM-CoA epimerase and MM-CoA mutase are both involved in fatty acid catabolism. MM-CoA epimerase catalyzes the rearrangement of (S)-methylmalonyl-CoA to the (R) form and uses a vitamin B12 cofactor (Overath et al. 1962). MM-CoA mutase induces the formation of adenosyl radical from AdoCbl cofactor and subsequently initiates a free-radical rearrangement of its substrate, (R)-methylmalonyl-CoA to succinyl-CoA - a key molecule of the citric acid cycle (Mancia et al. 1996). Methylmalonyl Co-A mutase-associated GTPase Cob (MeaB) is crucial for the proper functioning of methylmalonyl-CoA mutase (Takahashi-Iñiguez et al. 2017). Mutational analysis of this protein performed in *Methylobacterium* sp. showed an inability to convert methylmalonyl-CoA to succinyl-CoA caused by an inactive form of methylmalonyl-CoA mutase (Froese et al. 2010).

Kingdom Fungi comprises several lineages of non-Dikarya which, in the order of divergence, are classified into Chytrydiomycota and Blastocladiomycota grouping many aquatic organisms, fully terrestrial animal-related Zoopagomycotina, Entomophthoromycotina, Kickxellomycotina and plant/soil/dung-associated Mucoromycotina, Mortierellomycotina and Glomeromycotina. The remaining Dikarya are evolutionary youngest and best-studied fungal phyla and include Ascomycota and Basidiomycota. None of the aforementioned B12-related enzymes has been reported from fungi. Yet non-Dikarya, basal fungal lineages share multiple ancestral traits with animals and microbial eukaryotes. Here we show that all B12-dependent eukaryotic pathways are present in non-Dikarya fungi as well.

## Results

Our initial searches showed that only eight enzymes are uniquely present in B12-dependent organisms (Table 1). All of them have their homologs within early-diverging fungal lineages (Supplementary Table S1).

**Table 1.**
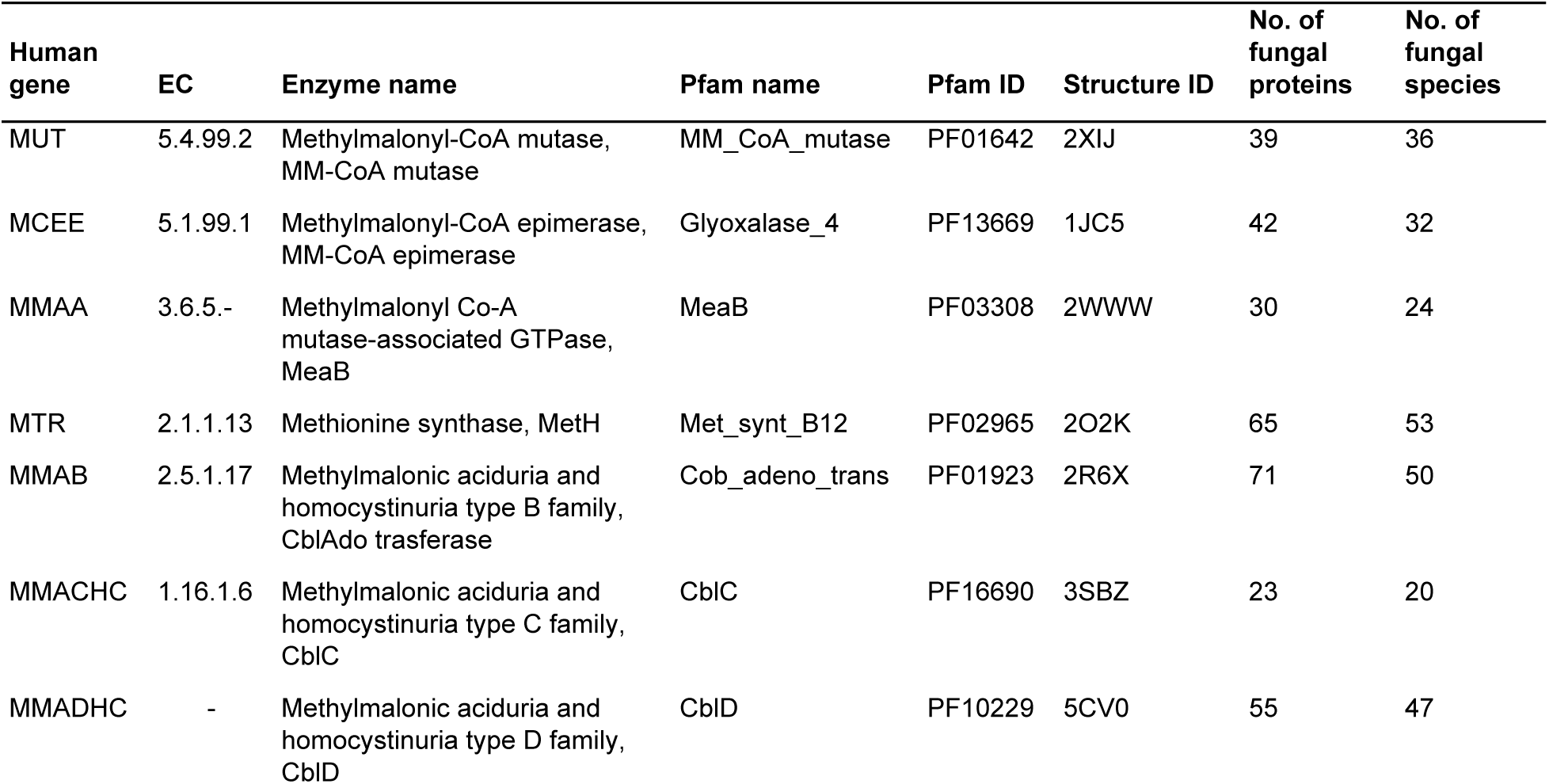
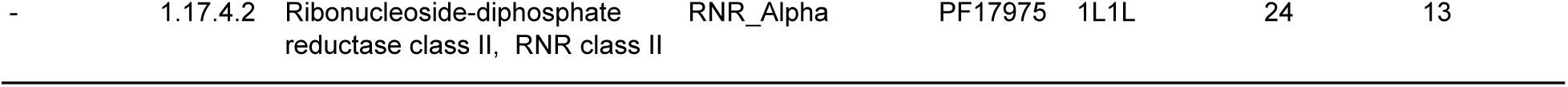
B12-specific enzymes used for the identification of B12-dependent pathways in fungal proteomes with the total number of homologs identified in this study.

### Distribution of B12 dependent enzymes in Fungi

Cobalamin-dependent enzymes were identified in 50 out of 59 analyzed non-Dikarya fungi proteomes (Table 1, see Supplementary Table S2 for detailed lists of all protein accessions). This dataset contains all genome derived protein predictions for all non-Dikarya isolates deposited in GenBank by October 2019, with representatives of all main lineages. Additionally, these enzymes were found in six Mucoromycota, one Blastocladiomycota, and two Cryptomycota representatives from NR database which were not included in the genomic dataset. Certain patterns in the distribution of cobalamin-related enzymes among non-Dikarya fungi can be seen at the phylum and lower taxa levels (Fig. 1, Supplementary Table S3). The whole set of studied enzymes is present in five non-Dikarya fungal proteomes, four of them belonging to the Glomeromycotina (Mucoromycota). The occurrence of cobalamin-related enzymes is common for all Mucoromycota species, but worth noting are the differences between Glomeromycotina, Mortierellomycotina and Mucoromycotina (the latter comprising saprotrophic Mucorales, Umbelopsidales and plant symbionts Endogonales). In Mucorales only three families of cobalamin-dependent enzymes are conserved (CblD, MetH, and CblAdo transferase). For other Mortierallomycotina and Endogonales, it is common to retain four or more of the analyzed protein families. The whole set of enzymes can be found also in Blastocladiomycota. Other taxa with high occurrence of cobalamin-dependent enzyme families are the animal-related Entomophthoromycotina, Kickxellomycotina, and Zoopagomycotina. All of them have homologs from four up to seven families.

**Figure 1.**
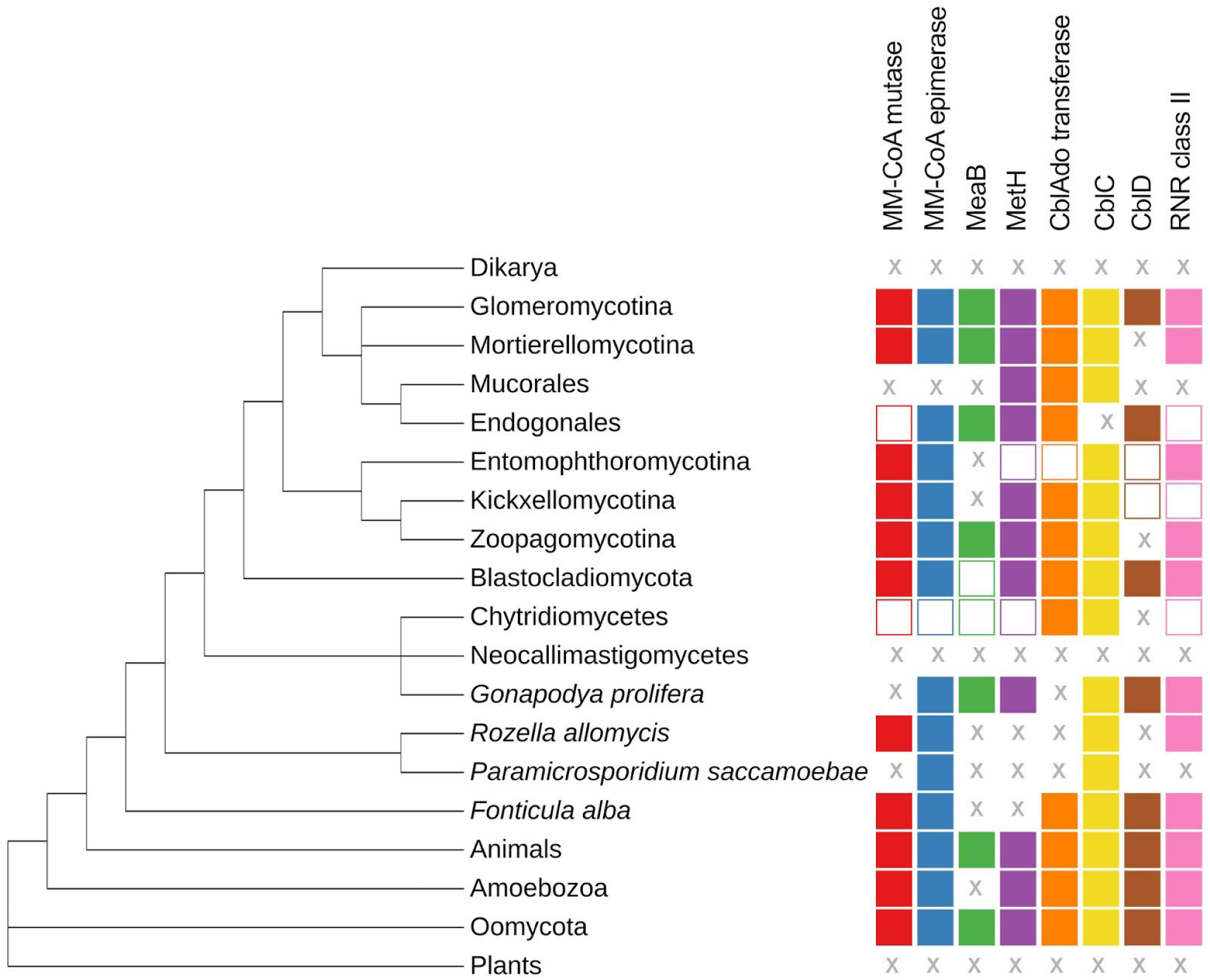
Distribution of B12-dependent protein families on a dendrogram showing a part of the eukaryotic tree of life, the schematic tree is based on (Davis et al. 2019, (Spatafora et al. 2016) for fungi and on (Burki et al. 2020) for remaining lineages. For each taxon, symbols on the right represent B12-dependent enzymes found in their proteome. X symbol means no identified homologs of the enzyme in the whole taxon; empty shape refers to the occurrence of the enzyme in less than half of studied representatives, filled shape means that half or more representatives have the enzyme in their proteomes.

Nine of the analyzed proteomes do not contain any of the studied enzymes. All these proteomes belong to Chytridiomycota. Chytridiomycetes and *Ganopodya prolifera* (Monoblephariodomycetes) have a wide range of enzymes occurrence - from zero in Caulochydriales, Rhizophydiales, and Blyttiomyces to six in Chytridiales and Monoblepharidales. Neocallimastigomycetes stand out especially here - none of the analyzed four proteomes from this taxon had any homologs of the cobalamin-related proteins family.

### Pathway conservation

Cobalamin-dependent enzymes play roles in three pathways associated with RNR class II, MetH, and MM-CoA mutase. Obtained results suggest that among early-diverging fungi there is a tendency to conserve the key enzymes rather than whole pathways. This is especially true in Mucorales which retained only CblAdo transferase and part of MetH pathways. In other non-Dikarya fungi, MM-CoA mutase associated pathway is also well conserved. The B12-dependent ribonucleotide reductase is least conserved but this might be associated with the presence of different RNR classes.

In order to ensure that all housekeeping functions provided by RNR class II, MM-CoA mutase, and MetH pathways are maintained present in all of the studied isolates, even those devoid of B12-dependent enzymes, we searched for cobalamin-independent alternatives, namely, MetE in place of MetH (González et al. 1996), RNR class I instead of class II (Jordan and Reichard 1998), and methylcitrate cycle (MCC) for MM-CoA mutase pathway (Dubey et al. 2013). We found that all these enzymes involved in B12-independent metabolic tracks are present in non-Dikarya fungi.

Since most of the identified homologs of eight B12-dependent enzymes are annotated as hypothetical unknown proteins without experimental characterization, we performed tblastn searches on them against the EST database. This served as intermediate evidence that the predicted B12-related proteins in non-Dikarya fungi originate from active genes. Tblastn search results allow also to expect that genes encoding all identified proteins will be expressed.

### Phylogenetic analysis

To trace the evolution of the studied proteins, phylogenetic trees for each of the eight protein families were inferred using Bayesian (BA) and Maximum Likelihood (ML) approaches (Supplementary Table S4, **Supplementary Figures**), except for MeaB and CblAdo transferase (with highest numbers of identified homologs) where BA analyses did not converge to a reliable level of the standard deviation of split frequencies. We noticed single bacterial sequences misannotated as fungal due to likely bacterial contamination of the fungal DNA samples. We also noticed single fungal sequences grouping within their bacterial relatives. In most cases these were proteins homologous to our enzyme yet with other function, e.g. MeaB is similar to other GTPases (KAA6408927.1). Non-Dikarya fungal sequences rarely grouped with bacterial sequences with the exception of MM-CoA mutase from *Syncephalis pseudoplumigaleata* (RKP28319.1, RKP28318.1) which displayed a very high sequence identity reaching 100% with *Afipia* alphaproteobacteria which might indicate sample contamination. Notably, Dikarya sequences did not enter Eukaryotic clades for the analyzed enzymes in contrast to their non-Dikarya homologs.

Exclusively fungal clades can be observed in five protein families (Fig. 3). For the other three enzymes, there are clades composed mostly of fungal homologs and ones belonging to other eukaryotic microorganisms (Holozoa, Amoebozoa, and SAR). Observed topologies in the eukaryotic part of the trees generally are congruent with the species tree. Interestingly, in two cases (CblAdo transferase and RNR class II) Oomycota and Fungi clades are sisters to each other. Sequences identity of randomly chosen homologs is ∼63% for RNR class II (ETI40368.1 and KNE69215.1) and ∼52% for CblAdo transferase (XP_002997018.1 and KNE71581.1).

**Figure 2.**
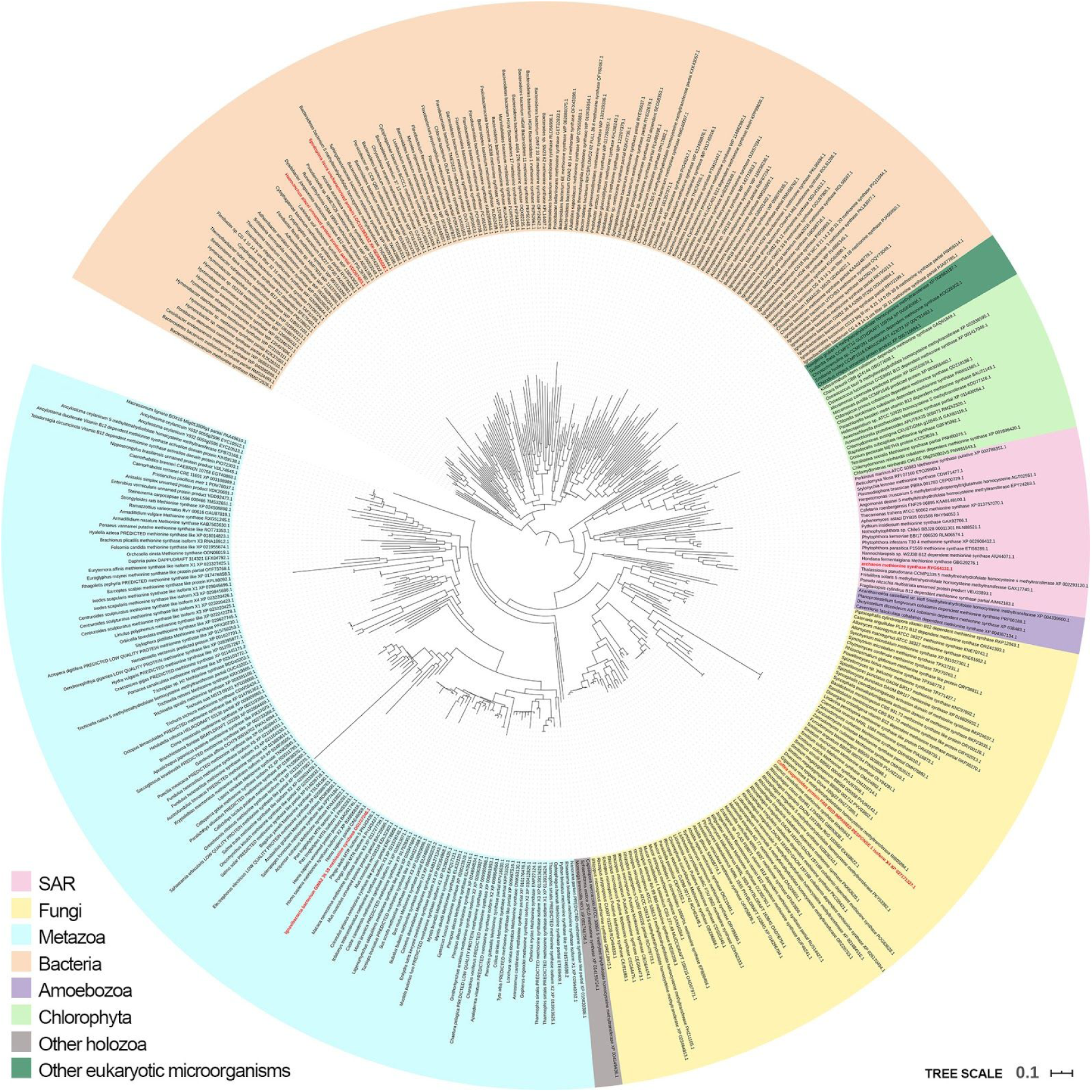
Phylogenetic tree of methionine synthase MetH homologs. The tree was built based on 72 sequences from non-Dikaryal proteomes, aligned with their homologues from NCBI non-redundant database 291 (Methods). Sequences marked with red labels do not belong to organisms to which they were assigned.

**Figure 3.**
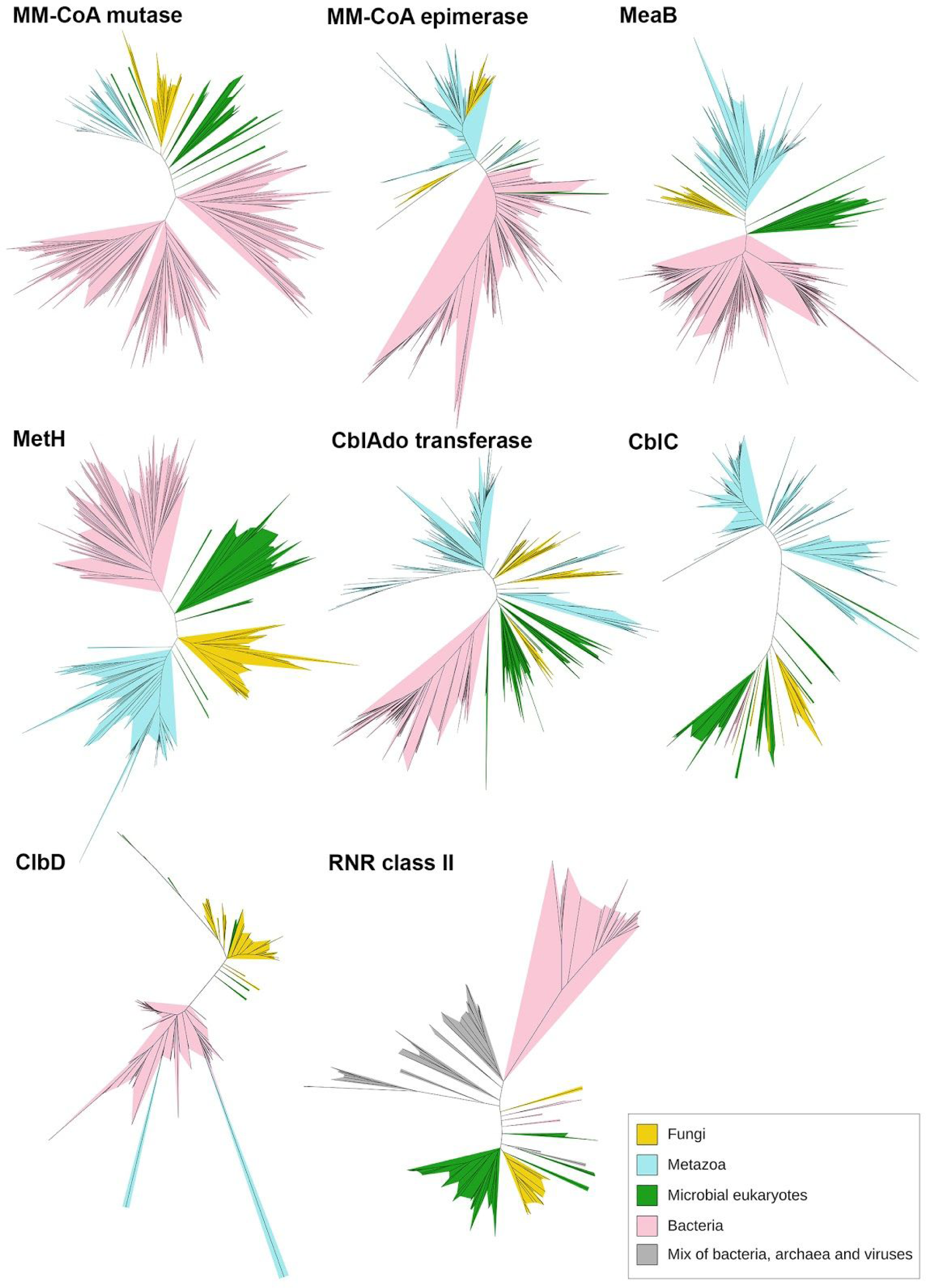
Unrooted ML trees of eight B12-related protein family representatives.

In five out of eight trees, fungal sequences form a monophyletic clade. For MeaB, CblD, MetH, MM-CoA mutase, and RNR class II, fungal sequences form a sister clade to *Fonticula alba* (Holomycota), the closest relative of fungi belonging to Nucleariida. This pattern was observed for class II with the following score of aLTR support of the fungal clade: MeaB - 1.00, CblD - 1.00, MetH - 1.00, MM-CoA mutase - 1.00, and RNR class II - 0.98. In the case of CblAdo transferase, CblC and MM-CoA epimerase fungal sequences group together with either ancient Metazoa representatives (CblAdo transferase) or with microbial eukaryotes from Holomycota, Amoebozoa and SAR groups (MM-CoA epimerase). Importantly, non-Dikarya fungal sequences are always sister to other Eukaryotic sequences which rules out bacterial contamination. Sequences from model organisms belonging to diverse lineages of microbial eukaryotes, not only Opisthokonta, were represented in these clades, including representatives from *Polysphondylium pallidum* and *Dictyostelium spp.* (Amebozoa), *Thecamonas trahens* (Apusozoa), *Chlamydomonas reinhardtii* (Chlorophyta), *Stentor coeruleus* (Alveolata), *Emiliania huxleyi* (Haptophyta), *Thalassiosira pseudonana* and *Blastocystis spp.* (Heterokonta), *Naegleria gruberi* and *Euglena gracilis* (Excavata).

## Methods

59 predicted fungal proteomes were downloaded from NCBI in October 2019 (Sayers et al. 2020) (Supplementary Table S1). Next, a pfam_scan.pl (default settings) (Mistry et al. 2007) search of all protein sequences against a library of Pfam HMMs was performed. We recognized the occurrence of domains from eight vitamin B12-dependent enzymes in non-Dikarya fungi proteomes (Table 1). To expand our dataset NR database was searched for homologs of those non-Dikarya fungal B12-dependent proteins and additionally for homologs of proteins from model eukaryotic organisms with known B12 dependent enzymes (*Homo sapiens, Dictyostelium discoideum, Fonticula alba, Phytophthora infestans*) using PSI-BLAST (evalue=0.001, num_iterations=3) (Altschul et al. 1997). The dataset was unified and clustered with CD-HIT (n=4, c=0.7, aS=0.95, aL=0.95), all fungal hits were retained regardless of their sequence similarity. To get only homologs of a protein of our interest, there was a need to discard homologs from related protein families. To do this we visualized protein pairwise similarity using CLANS (Frickey and Lupas 2004; Mistry et al. 2007) and selected separated groups of sequences.

In the next step, sequences were aligned using local iterative mode in Mafft v. 3.7 (localpair, maxiterate=100) (Katoh et al. 2002). The alignment was additionally cleared manually from potential inactive homologs. All sequences that showed a lack of amino-acids crucial for enzyme activity or substitution of them with amino-acids that are not able to maintain enzyme activity, were discarded from the set.

All alignments were trimmed with TrimAl (model=gappyout) (Capella-Gutiérrez et al. 2009) to remove poorly conserved regions. Then, by using ProtTest (all-matrices, all-distributions) (Abascal et al. 2005; Capella-Gutiérrez et al. 2009), we appointed the best amino-acid substitution models based on Akaike Information Criterion AIC. Phylogenetic trees were built using LG model for each of the B12 metabolism-related enzymes with Bayesian (BA) and Maximum likelihood (ML) approaches using MrBayes (Huelsenbeck and Ronquist 2001) and PhyML (Guindon et al. 2010) respectively. ML trees were estimated with a gamma distribution of rates between sites (four categories and alpha parameter estimated by PhyML) and aLRT Chi2-based parametric branch supports. In the course of BA inference, four Markov chains were run for 3 runs from random starting trees for 107 generations, and trees were sampled every 2,5×102 generations. The first one-fourth of generations were discarded as burn-in. Then, we used the remaining samples to calculate the tree of maximum clade credibility.

Expression of representatives of each of protein sets (three randomly chosen homologs from each family) was confirmed by tblastn (Altschul et al. 1990) (default settings) searches against the EST database at NCBI website.

## Discussion

Our discoveries contradict the current opinion that fungi neither synthesize nor use cobalamin (Duda et al. 1957; Jah et al. 2002) and do not have cobalt at all (Zhang et al. 2019). This claim remains true for Dikarya, but we demonstrate that the older fungal lineages do have proteins that either process or use cobalamin as a cofactor.

In Eukaryotes three main metabolic pathways use cobalamin - RNR class II, MM-CoA mutase and MetH pathways. Functions provided by these pathways are needed for the independent functioning of a living cell and can be lost in parasites (Zhang et al. 2009). Transport and trafficking of cobalamin in the cell is described in animals but homologs of the proteins responsible for the cobalamin transport e.g. LMBR1-like membrane protein transporters have a universal distribution in the Opisthokonta. Many of the enzymes involved in the MM-CoA mutase and MetH pathways, like mevalonate kinase and methionine synthase reductase, respectively, are conserved independently of B12 usage.

We found traces of all of these pathways among the whole fungal tree of life, except Dikarya lineage. The distribution of the genes encoding the above-mentioned enzymes is not uniform across the analyzed organisms. For non-Dikarya fungi, it is common to either have two out of three pathways or to have them incomplete.

Only Glomeromycotina and Blastocladiomycota have all three complete B12-dependent pathways. These two taxonomic groups are evolutionary and ecologically distant, they share only a few characteristics, including big genomes with a high number of genes. The latter may be a highlight of relaxed pressure on genome compactness.

The least conserved among fungal lineages is the RNR class II pathway. However, organisms missing this class use cobalamin-independent RNR class I, which is common in animals. It is worth noticing that organisms are not limited to having only one class of RNRs at once (Jordan and Reichard 1998). Cobalamin-dependent RNR class II appears mostly in bacteria and, according to our results, also in non-Dikarya fungi and Oomycota. Additionally, RNR class II homologs from these groups can be found in sister clades (this is also true for CblAdo transferase homologs). This may suggest an ancient horizontal gene transfer between Oomycota and fungi resulting in nonidentical but highly similar sequences. This is yet another parallel molecular trait that groups fungi and other filamentous fungi-like organisms together, next to similarities in weaponry to attack plants (Latijnhouwers et al. 2003), the evolution of the nitrate assimilation pathway (Ocaña-Pallarès et al. 2019), and the role of horizontal gene transfer (Rosewich and Kistler 2000; Soanes and Richards 2014). This trait is exquisitely interesting because it is shared by eukaryotic microorganisms but is absent from big multicellular forms.

The best-conserved pathway in non-Dikarya fungi - MetH - can be substituted with a cobalamin-independent enzyme called MetE (González et al. 1996). We checked if this enzyme variant also can occur in non-Dikarya fungi proteomes. MetE is present in all non-Dikarya fungi phyla, even in Neocallimastigomycetes, which do not have any other cobalamin-dependent or independent alternatives of studied pathways. For some of the non-Dikarya fungi, lack of CblC protein can be observed. We did not look for substitutes for this protein, because the cooperation of CblC and CblD in the MetH pathway was described only for animals - outside this group, the exact function of CblD protein is not documented, and perhaps in other organisms, CblC is sufficient to perform its function by itself. One might speculate that other proteins are recruited to catalyze decyanation of cyanocobalamin and dealkylation of alkylcobalamins in non-animal organisms.

MM-CoA mutase pathway is more or less conserved among early-diverging fungal lineages. Interestingly, all Mucorales members lack all three B12-dependent enzymes of that pathway. We checked for alternatives for this metabolism track and it turned up to be more complex than in the other two cases. In Dikarya propionate metabolism is carried out in the methylcitrate cycle (MCC). Three key enzymes for this track are methylcitrate synthase (MCS), methylcitrate dehydrogenase (MCD), and methylisocitrate lyase (MCL) (Dubey et al. 2013). All of them are present in Dikarya and, interestingly also in Choanoflagellida and Metazoa, but not in early-diverging fungal and other ancient lineages like Ichthyosporea and in Fonticula. MCS and MCL are conserved as well in old fungal phyla as in Dikarya, but that does not apply to MCD. Following information about the MCC gene cluster (Santos et al. 2020) genomic context of this pathway was checked for *Batrachochytrium* and *Mucor* representatives showing no synteny. Moreover, no candidate dehydrogenases were found upstream or downstream of MCS and MCL genes. We assume that the function of MCD can be taken over by other dehydrogenases.

According to our results, we can speculate the best-conserved elements of cobalamin-dependent pathways are key enzymes. For example, in the MetH pathway, the best-conserved element is MetH protein. On the contrary, it is quite common to lose CblC and CblD proteins from the proteome (Supplementary Table S1). The question is why in some organisms only the main part of pathways is conserved and how it is possible for these pathways to work without helper protein. We speculate that our results may be biased towards the main enzymes because they are well-known, especially have a well-known active site what allows for more rational data curation. Because our selection of potentially active homologs heavily relied on identified active site residues, it could have resulted in an underestimate of helper protein identification. Additionally, the MetH pathway is well described only in animal metabolism, so we cannot be sure about the role of CblC and CblD in fungal metabolism and about the necessity of having these proteins. On the other hand, the best-conserved protein in the MM-CoA mutase pathway is CblAdo transferase. For Mucorales, it is common to have only this one protein from the whole MM-CoA mutase pathway. It is worth noticing that this protein is responsible for synthesizing AdoCbl cofactor for MM-CoA mutase which is the only protein in fungal metabolism that is known to require the AdoCbl cofactor. The question is why in Mucorales proteomes there is still pressure to conserve CblAdo transferase while it is common to lose MM-CoA mutase.

Literature suggests that host-associated organisms have a tendency for the loss of cobalt utilization pathways (Zhang et al. 2009). Our results suggest that non-Dikarya fungi comply with this assumption. Chytridiomycota phylum combines amphibian parasites *Batrachochytrium* sp. and herbivorous mammals symbionts from class *Neocallimastigomycetes.* For these organisms, no cobalamin-dependent enzyme was found. These organisms may obtain the required resources from the host. However, our observations for plant-associated fungi are different. Mycorrhizal fungi from Glomeromycotina and Endogonales, despite maintaining extensive symbiotic relationships with 80% of plant species (Smith and Read 2010), retain well-conserved cobalamin-dependent pathways.. It is possible that plant-associated non-Dikarya fungi kept these pathways simply because plant metabolism lacks cobalamin. The difference between plant and animal associated fungi may be a consequence of different pressures in such diverse ecological niches. Generally, parasites and obligate symbionts are biotrophs characterized by reduced genome size. However, in the case of mycorrhizal fungi, for a yet unknown reason, the pressure to reduce the genome seems to be relaxed.

The question that still remains is what is the source of cobalamin for fungi. We speculate that fungi are able to accumulate B12 acquired from bacterial sources. B12 cofactor supply for at least some of the fungi with B12-dependent enzymes may be mediated by endohyphal bacteria with an intact B12 synthesis pathway. All crucial components of the B12 *de novo* synthesis pathway required for such a relationship were found in the case of symbiosis between Glomeromycotina fungus *Gigaspora margarita* and β-proteobacterium *Candidatus Glomeribacter gigasporarum* (Ghignone et al. 2012).

During the analysis of obtained results, we tried to understand the evolution of cobalamin-dependent metabolic pathways among kingdom Fungi. To widen the picture we checked studied proteomes for cobalamin-independent alternative metabolic pathways, and we confirmed their occurrence. Based on current knowledge we hypothesized that B12-dependent pathways are replaced by B12-independent alternatives in course of the evolution, and finally disappear in Dikarya lineages. In fact, the ability to utilize cobalamin is either retained or lost independently from the time of phyla divergence. Surprisingly there is a much stronger correlation between the preservation of this ability and fungal ecology. As we observed, cobalamin-dependent pathways are more common in fungi associated with plants, than in species associated with animals and living as soil saprophytes.

Our discovery challenges the current view that fungi can neither synthesize nor utilize cobalamin. We proved that non-Dikarya fungal proteomes contain three metabolic pathways utilizing vitamin B12. We speculate these organisms have the possibility to accumulate cobalamin. Our discoveries may open the way for the selection of B12 over accumulating strains of food fermenting fungi without the need for genetic material manipulation.

### B12 enzymes in other eukaryotes

We also confirmed the occurrence of B12 related enzymes in other Holomycota taxa like Cryptomycota (Rozellida and Microsporidia) and Fonticulida. These organisms retain a maximum of only six out of the eight enzymes, but it is worth noting they are not independent, free-living organisms.

Studied enzymes are also present in Amoebozoa and Oomycota. Some other species from the SAR supergroup, to which Oomycota belongs (Burki et al. 2020), are known to have cobalamin-dependent methionine synthase (Boudouresque 2015). The matter is not clear about B12 utilization in Amoebozoa. There is contradictory information on the necessity to supplement the culture of *Dictyostelium discoideum* with that vitamin (Stephan et al. 2003). In addition, class II RNR has been observed in *D. discoideum* previously (Crona et al. 2013). B12-dependent enzymes are encountered also in green algae (Chlorophyta), red algae (Rhodophyta) (Croft et al. 2005; Thi Vu et al. 2013)) and Excavata (Helliwell et al. 2016). Green algae are known to acquire vitamin B12 through a symbiotic relationship with bacteria (Croft et al. 2005; Thi Vu et al. 2013).

Taken together, B12 dependence seems to be a widely distributed trait in Eukaryotes and was likely present in the last common ancestor of Eukaryotes. Several multicellular lineages including vascular plants and Dikarya developed B12-independent alternative pathways and, eventually, lost the B12 metabolism completely.

## Supplementary Legends

Supplementary Table S1 Table with numbers of each eight cobalamin-dependent enzymes homologs in all studied non-Dikarya fungi.

Supplementary Table S2 Accessions of all identified protein homologs.

Supplementary Table S3 PMIDs of sources of information about all studied genomes.

Supplementary Dataset DS1 File with phylogenetic trees of B12-dependent enzymes in non-Dikarya fungi in Newick format. Contains ML trees with all found homologs (full) and chosen subtree and BA subtree of all studied enzymes.

## Acknowledgements

We thank Anna Karnkowska, Julia Pawłowska, Krzysztof Pawłowski and Marcin Grynberg for their insight and comments about the manuscript. This work was supported by the Polish National Science Centre (grant 2017/25/B/NZ2/01880 to AM).

## Data Availability Statement

The analyses are based on publicly available sequences. All identifiers of analysed proteins are listed in the Supplementary Materials. All trees built from those sequences are also listed in the Supplementary Materials.

## Author Contributions

A.M. designed the study, M.O. and A.M. prepared the data set and performed sequence analyses and M.O., K.S. and A.M. interpreted the data and wrote the manuscript.

